# *Plasmodium falciparum* infection of human erythroblasts induces transcriptional changes associated with dyserythropoiesis

**DOI:** 10.1101/2023.04.23.538003

**Authors:** Tamar P. Feldman, Yana Ryan, Elizabeth S. Egan

## Abstract

During development down the erythroid lineage, hematopoietic stem cells undergo dramatic changes to cellular morphology and function in response to a complex and tightly regulated program of gene expression. In malaria infection, *Plasmodium spp*. parasites accumulate in the bone marrow parenchyma, and emerging evidence suggests erythroblastic islands are a protective site for parasite development into gametocytes. While it has been observed that *Plasmodium falciparum* infection of late-stage erythroblasts can delay terminal erythroid differentiation and enucleation, the mechanism(s) underlying this phenomenon are unknown. Here, we apply RNA-seq after fluorescence-activated cell sorting (FACS) of infected erythroblasts to identify transcriptional responses to direct and indirect interaction with *Plasmodium falciparum*. Four developmental stages of erythroid cells were analyzed: proerythroblast, basophilic erythroblast, polychromatic erythroblast, and orthochromatic erythroblast. We found extensive transcriptional changes in infected erythroblasts compared to uninfected cells in the same culture, including dysregulation of genes involved in erythroid proliferation and developmental processes. Whereas some indicators of cellular oxidative and proteotoxic stress were common across all stages of erythropoiesis, many responses were specific to cellular processes associated with developmental stage. Together, our results evidence multiple possible avenues by which parasite infection can induce dyserythropoiesis at specific points along the erythroid continuum, advancing our understanding of the molecular determinants of malaria anemia.

**Key Points:**

- Erythroblasts at different stages of differentiation have distinct responses to infection by *Plasmodium falciparum*.
- *P. falciparum* infection of erythroblasts alters expression of genes related to oxidative and proteotoxic stress and erythroid development.

## Introduction

Malaria is a mosquito-borne infectious disease caused by parasites of the genus *Plasmodium*. Recent data estimates 593,000 fatalities due to malaria in the year 2021, the large majority of which occurred among young children and pregnant women in sub-Saharan Africa.^1^ Whereas five different species of *Plasmodium* parasites can cause disease in humans, most malaria deaths are caused by *Plasmodium falciparum*.^2^ During infection, parasites undergo rounds of intracellular replication in erythrocytes (red blood cells) and sequester in the microvasculature and deep tissues, including sites of hematopoiesis in the bone marrow.^2–4^ Consequently, malaria infection is often complicated by severe anemia related to increased destruction or clearance of erythrocytes.^5–8^ This adverse effect can be further compounded by failure of the bone marrow to adequately produce and release new erythrocytes into circulation.^9–12^

Bone marrow dyserythropoiesis remains a poorly understood aspect of the pathogenesis of malarial anemia, but has been observed in acute, severe disease and in chronic infections common in regions of high transmission. Bone marrow aspirates of malaria patients with severe anemia demonstrate morphological abnormalities in developing erythroblasts, including nuclear fragmentation, multinucleate cells, and inter-cytoplasmic bridges.^10, 12^ Asexual and sexual stage *Plasmodium spp.* parasites have been detected in the bone marrow of patients with malaria, including adjacent to erythroblastic islands.^13–15^ Indeed, the bone marrow is now recognized as a protective niche for developing *P. falciparum* gametocytes, and recent work suggests that sexual commitment is influenced by the nutrient state of the host cells.^13, 14, 16, 17^ Similar findings have also been reported in rodent malaria models.^18, 19^ Hemozoin, an iron biocrystal produced by parasite digestion of hemoglobin, is also present in the bone marrow during malaria infection and has been implicated in the pathogenesis of malaria anemia.^10, 13, 14, 20–22^ Together, these studies raise the hypothesis that the interaction of developing erythroid precursor cells with *Plasmodium* parasites and metabolites in the hematopoietic niche may contribute to disordered erythropoiesis.

Advances in culture systems for *ex-vivo* erythropoiesis enable study of host-parasite interactions during erythroid differentiation.^20^ Using specialized growth media, primary human hematopoietic stem/progenitor cells (HSPCs) can be directed down the erythroid lineage to produce enucleated reticulocytes.^23–25^ Along the way, erythroblasts pass through the canonical stages of terminal differentiation: proerythroblast (ProE), basophilic erythroblast (BasoE), polychromatic erythroblast (PolyE), and orthochromatic erythroblast (OrthoE). Efforts to characterize the dynamic changes in expression of membrane surface proteins during terminal erythroid differentiation have identified markers for each erythroblast stage, facilitating stage-specific isolation for functional analyses.^26–28^

Primary human erythroblasts cultured *ex-vivo* are susceptible to infection with *P. falciparum* as early as the BasoE stage, but the impact of parasite infection on erythroid differentiation is poorly understood.^17, 29^ Increased reactive oxygen species (ROS) and reduced enucleation have been observed in late-stage erythroblasts harboring gametocytes, and hemozoin has been shown to inhibit growth of primary erythroblasts and to induce cell cycle changes in K562, an erythroleukemic cell line.^21, 30–32^ Transcriptional profiling by microarray has shown that co-culture of *P. falciparum* with PolyE and OrthoE may perturb expression of host gene pathways related to metabolism and protein chaperones, though the bulk nature of these studies limit firm conclusions.^33^ Hemozoin has also been shown to alter transcription of genes involved in apoptosis in late-stage erythroblasts.^34^ Overall, the host and parasite factors that underlie these effects are largely unknown.

To bridge the gap in knowledge, we leveraged an *ex-vivo* erythropoiesis culture system for primary human HSPCs to develop a strategy for transcriptomic profiling of infected and uninfected (bystander) erythroblasts at distinct stages of terminal erythroid differentiation. We used fluorescence activated cell sorting (FACS) followed by RNA-seq to investigate the host transcriptional response to infection or exposure to *P. falciparum* in ProE, BasoE, PolyE, and OrthoE. Our results demonstrate shared and distinct host cell responses to *P. falciparum* according to the differentiation state of the erythroblast and suggest that direct infection alters the expression of genes that are critical for progression of healthy erythropoiesis.

## Methods

### Primary human CD34^+^ HSPC culture

A three-stage protocol was used to differentiate human bone marrow CD34^+^ cells (STEMCELL Technologies) down the erythroid lineage as previously described.^37^ Briefly, CD34^+^ cells were cultured in IMDM-based medium with plasma and supplemented with IL-3, hydrocortisone, Stem Cell Factor (SCF), and erythropoietin (EPO). On day 7 of differentiation, IL-3 and hydrocortisone were removed from the supplementation to induce terminal differentiation. From day 11 until the end of culture, cells were switched to medium supplemented with only EPO. Differentiation was monitored by cytospins stained with May-Grünwald and Giemsa, and visualized by light microscopy.

### P. falciparum culture

*P. falciparum* parasites were grown in a RPMI 1640-based medium (Sigma) supplemented with 25 mM HEPES (Gibco), 50 mg/L hypoxanthine (Sigma), and 0.5% Albumax (Invitrogen) at 37°C in 1% O_2_ and 5% CO_2_ with gentle agitation. Parasites were maintained at 2% hematocrit in de-identified human erythrocytes obtained from Stanford Blood Center. Erythroblast infections used *P. falciparum* strain D10-pfPHG (provided by Dave Richard).^35^ D10-pfPHG is a laboratory-adapted strain expressing cytosolic GFP under the PfHsp86 promoter.

## Infection of primary human erythroblasts

Erythroblasts were removed from the parent culture and plated at 1 ×10^6^/mL in IMDM-based media supplemented with Albumax, hypoxanthine and cytokines corresponding to the day of differentiation. Parasites were synchronized to a 1-2 hour window one cycle prior to infection. Schizont-stage parasites were purified using an LS magnetic column (Miltenyi) and mixed with erythroblasts at an MOI of 5 (day 7) or 3 (day 13) unless otherwise specified. Static co-cultures were incubated for 24h at 37°C in 1% O_2_ and 5% CO_2_.

### Flow cytometry and cell sorting

Cells were analyzed and sorted on a FACSAriaII (BD Biosciences). Erythroblasts were staged using CD235a-APC eFluor780 (Thermofisher), CD233-PE (IBGRL), and CD49d-APC (Miltenyi). Viability was determined using Calcein-Violet 450AM. Infected erythroblasts were identified by GFP expression from the parasite. Cells were sorted directly into buffer RLT (Qiagen) for RNA extraction or into FBS for downstream microscopy.

### Library preparation and sequencing

Total RNA was extracted from sorted cells using the RNeasy Micro kit (Qiagen) and checked for quality and concentration using the Agilent Eukaryotic RNA 6000 Pico kit run on an Agilent 2100 Bioanalyzer at the Stanford Protein and Nucleic Acid Facility. Paired-end libraries were prepared using the Tecan Trio RNA-seq kit for low-input samples with automation on the Agilent BRAVO. Custom AnyDeplete probes (Tecan) were added against the mitochondrial genome, *HBB*, *EEFA1*, *MALAT1*, and *HSP90aa1* to improve sequencing depth of low and moderately expressed transcripts. Pooled libraries were sequenced on an Illumina NovaSeq 6000 S1 flowcell using a 200 cycle SBS kit.

### Quantification and statistical analysis

Reads were aligned to a concatenation of the *Homo sapiens* and *P. falciparum* genomes. Subsequent analysis of human and parasite gene expression was conducted separately. Differential gene expression analysis was performed using DESeq2. Gene list enrichment analysis was conducted using EnrichR and the MSigDB Hallmark 2020 gene sets. Comparison of the parasite transcriptome in erythroblast infection to the published transcriptome of the asexual stage parasites^36^ was conducted by Spearman’s correlation on gene lists ranked by expression.

Further information on methods is available in the *Online Supplemental Appendix*.

### Data sharing statement

All data generated or analyzed during this study be found in the data supplements available with the online version of this article. Raw sequencing data can be found in the NCBI Gene Expression Omnibus (GEO) database.

## Results

### P. falciparum infects erythroblasts at all stages of terminal erythropoiesis

To investigate infection of developing erythroblasts with *P. falciparum*, we differentiated primary CD34^+^ cells from human bone marrow down the erythroid lineage using our previously published approach for *ex-vivo* erythropoiesis (**Figure 1A**).^37^ We performed infections by incubating schizont-stage parasites in excess with erythroblasts at 7, 10, or 14 days of *ex-vivo* culture (**Figure 1B**). Ring-stage parasites were visible in the cytoplasm of BasoE, PolyE, and OrthoE on cytospins prepared after 18-20h of co-culture, but typically dark staining of the cytoplasm precluded observation of infection in ProE cells (**Figure 1C**). To measure parasitemia in the erythroblast cultures, we used flow cytometry following infection with a GFP-expressing parasite strain, D10-PfPHG.^35^ The percent of GFP^+^ erythroblasts was higher in the day 14 infection relative to day 10, consistent with prior reports of robust infection in late-stage erythroblasts (**Figure 1D**). Cell sorting was used to confirm that the GFP^+^ population represented infected erythroblasts (**Figure 1E**).

**Figure 1:**
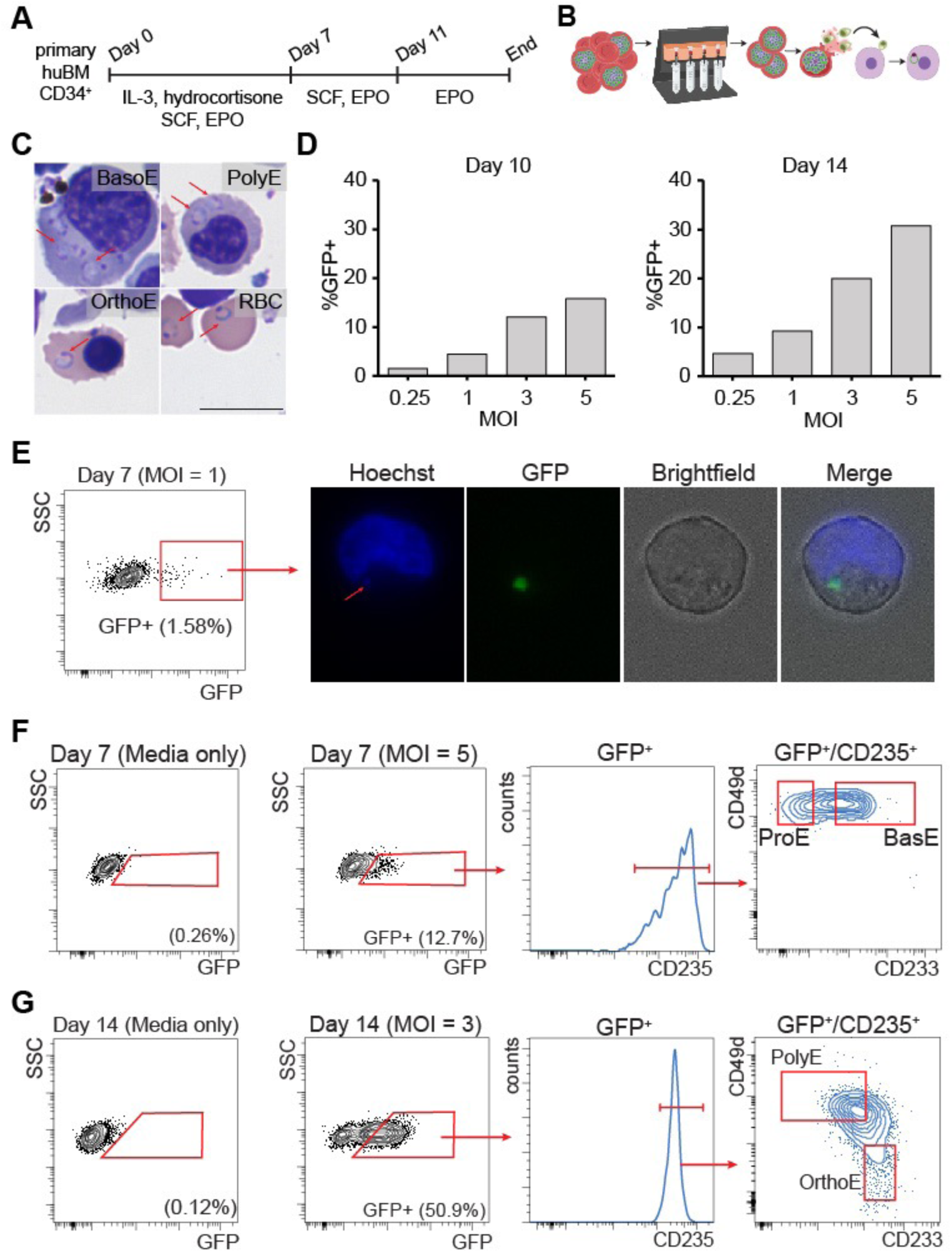
*P. falciparum* infects erythroblasts at all stages of terminal differentiation. (A) Schematic summarizing our primary cell culture system for ex vivo erythropoiesis. (B) Schematic depicting positive selection of late-stage *P. falciparum* by magnetic column prior to infection of erythroblast cultures. (C) May-Grünwald Giemsa staining showing ring-stage parasites in the cytoplasm of erythroid cells at multiple stages of terminal differentiation. Samples were collected for 18-20h post-mixing with schizonts. Scale bar represents 10μm. Arrows indicate ring-stage parasites. (D) Percentage of GFP^+^ erythroblasts measured by flow cytometry 18-20h after mixing with schizonts from a GFP-expressing parasite strain, D10-PfPHG. Erythroblasts were infected on day 10 (left) or day 14 (right) of differentiation. (E) Gating for FACS of GFP^+^ erythroblasts 18-20h after infection with D10-PfPHG at MOI = 1. Erythroblasts were infected on day 7 of differentiation. Fluorescence microscopy showing localization of GFP in an infected erythroblast sorted from the GFP^+^ population. Arrow indicates Hoechst staining overlapping GFP. (F) Flow cytometry analysis showing staging of infected, day 7 erythroblasts into proerythroblast (ProE) and basophilic erythroblast (BasoE). (G) Flow cytometry analysis showing staging of infected, day 14 erythroblasts into polychromatic erythroblast (PolyE) and orthochromatic erythroblast (OrthoE).

We next investigated the stage-specific susceptibility of human erythroblasts to infection with *P. falciparum*. We hypothesized that erythroblasts at all stages of terminal erythropoiesis are infectable by *P. falciparum*, although not all stages may support growth to the schizont stage. To identify erythroblast stages, we used a panel of antibodies targeting surface proteins that distinguishes between stages (anti-CD235a, anti-CD233, and anti-CD49d^26^), as confirmed by cytospin of sorted populations (**Figure S1**). When erythroblasts were infected at 7 days of differentiation, we detected GFP^+^ ProE and BasoE at 18-20 hours post-infection (hpi, time from addition of schizonts) (**Figure 1F**). Infection at 14 days of differentiation resulted in GFP^+^ PolyE and OrthoE at 18-20hpi (**Figure 1G**). Thus, parasites can establish infection in erythroblasts at all stages of terminal erythropoiesis.

### RNA-seq after FACS reveals host cell responses to P. falciparum infection of erythroblasts

To uncover stage-specific host cell responses to *P. falciparum* infection of erythroid precursors, we designed and implemented a FACS-based RNA-seq strategy that can disentangle the effect of erythroblast maturity on transcriptomic measurements and distinguish the response of infected cells from their uninfected neighbors (**Figure 2A**). Primary human CD34^+^ cells were induced to proliferate and differentiate down the erythroid lineage. At two timepoints of differentiation, day 7 and day 13, erythroblasts were removed from culture and plated with D10-PfPHG schizonts in six replicate wells that served as biological replicates. Erythroblasts mixed with media only were included as a reference for erythroid development in the absence of *P. falciparum*. A condition in which erythroblasts were incubated with dead, syringe-ruptured schizonts (SRS) was also included to identify host responses due to parasite debris and metabolites released upon rupture. Cells were harvested at 24hpi and stained with the panel of antibodies targeting surface proteins that distinguishes between erythroblast stages. Using FACS, we isolated ProE, BasoE, PolyE, and OrthoE erythroblasts from four different exposure categories: media-only (media), SRS-exposed (SRS), *P. falciparum*-infected (GFP^+^, infected), and *P. falciparum*-exposed but uninfected (GFP^-^, uninfected). After completion of the workflow, each population had at least four biological replicates for analysis (**Supplementary Datasets 1, 2**).

**Figure 2:**
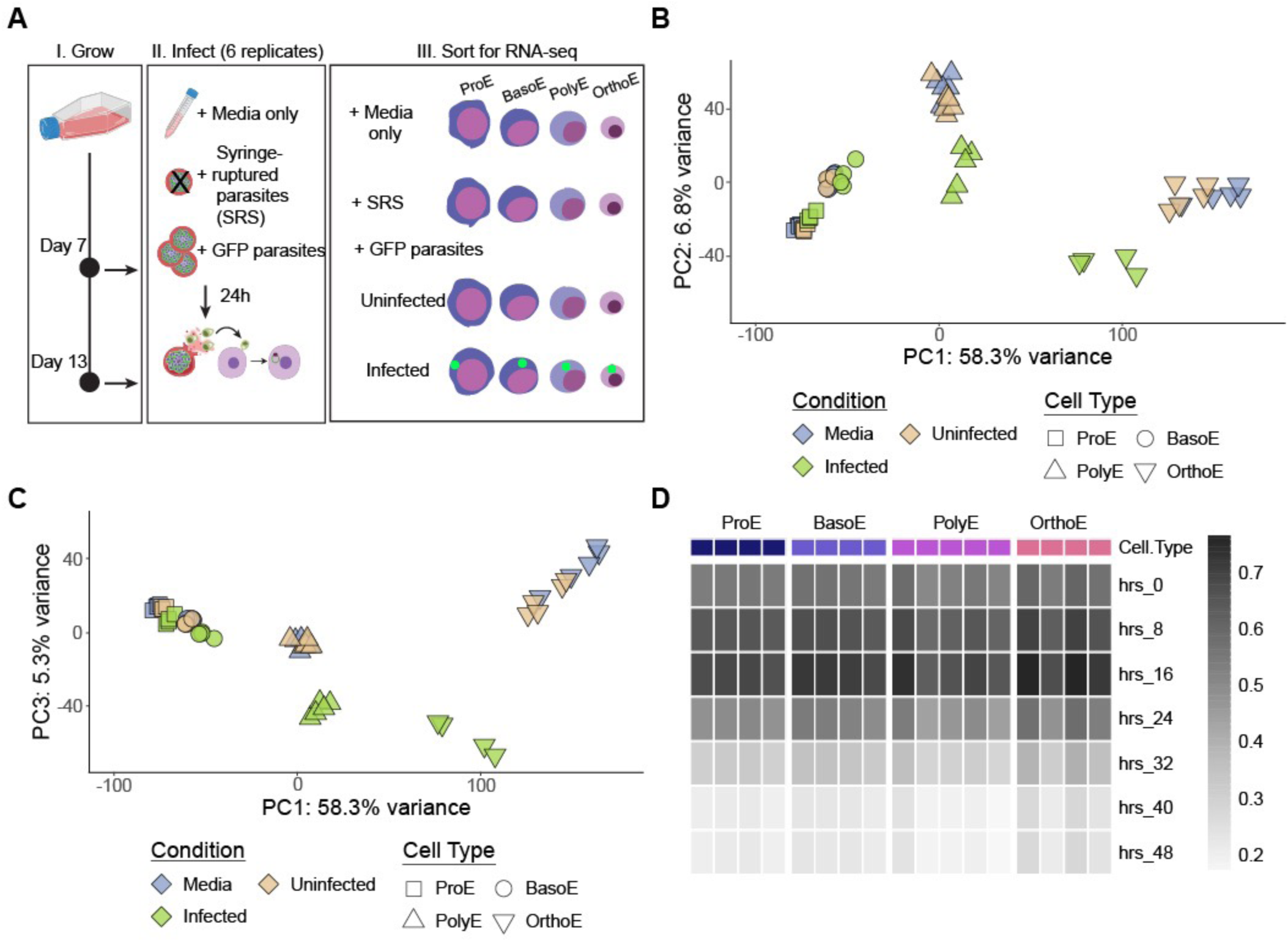
RNA-seq after FACS enables the study of host responses to *P. falciparum* at the terminal stages of erythroid differentiation. (A) Schematic of experimental design for characterizing transcriptional host responses to *P. falciparum* in specific erythroblast populations. Erythroblasts were removed from culture on day 7 or day 14 and mixed with GFP-expressing *P. falciparum* schizonts or an equivalent volume of media. After 24h, infected and uninfected erythroblast populations were collected by FACS for RNA extraction and transcriptomic analysis by RNA-seq. (B-C) Principal component analysis of RNA-seq dataset. Marker shape indicates erythroblast population. Color represents infection condition. D) Heatmap showing Spearman’s correlation between the parasite transcriptome in nucleated erythroblasts and the intraerythrocytic parasite transcriptome measured every eight hours as published by Otto et al.^36^ Colored annotations represent cell type, and each column represents a single replicate.

Principal component analysis (PCA) based only on human gene counts of each sample highlighted the trajectory of erythroid development (**Figure 2B, C**). Samples from the SRS condition grouped with uninfected and media-only conditions (**Figure S2**). In contrast, infected populations at each erythroblast stage appeared distinctly grouped from the other stage-matched populations based on host gene expression. From this preliminary analysis, we chose to focus our detailed analysis on the distinct host responses found in infected cells compared to uninfected cells and to the media-only condition. Notably, we found that the proportion of reads mapping to the parasite genome in infected populations increased as erythroblasts matured, consistent with the dynamic reduction in transcriptional activity known to occur in the final stages of terminal erythropoiesis (**Supplementary Table S1**). Comparison of our *P. falciparum* RNA-seq data with published data of *P. falciparum* gene expression during asexual growth in erythrocytes showed a strong correlation with the ring-stage of development and did not appear to differ by erythroblast stage (**Figure 2D)**.

### Infection with P. falciparum induces stage-specific changes in the host transcriptome at all stages of terminal erythropoiesis

To explore the host cell response to *P. falciparum* infection of the erythroid precursors, we applied differential gene expression analysis to pair-wise comparisons of media-only, uninfected, and infected populations at each erythroblast stage (**Supplementary Datasets 3 - 6**). As expected, the total number of detected human genes decreased over each consecutive stage of differentiation to reflect an increasingly specialized transcriptome (**Figure 3A-D**). We found that less than 4% of the transcriptome was differentially expressed between uninfected ProE (cells exposed to but not infected by *P. falciparum*) compared to media-only, and less than 2% at other erythroblast stages. These results suggest that the erythroblasts had minimal transcriptional responses when exposed to infected cells and parasite debris, at least at the 24-hour timepoints used in these experiments.

**Figure 3:**
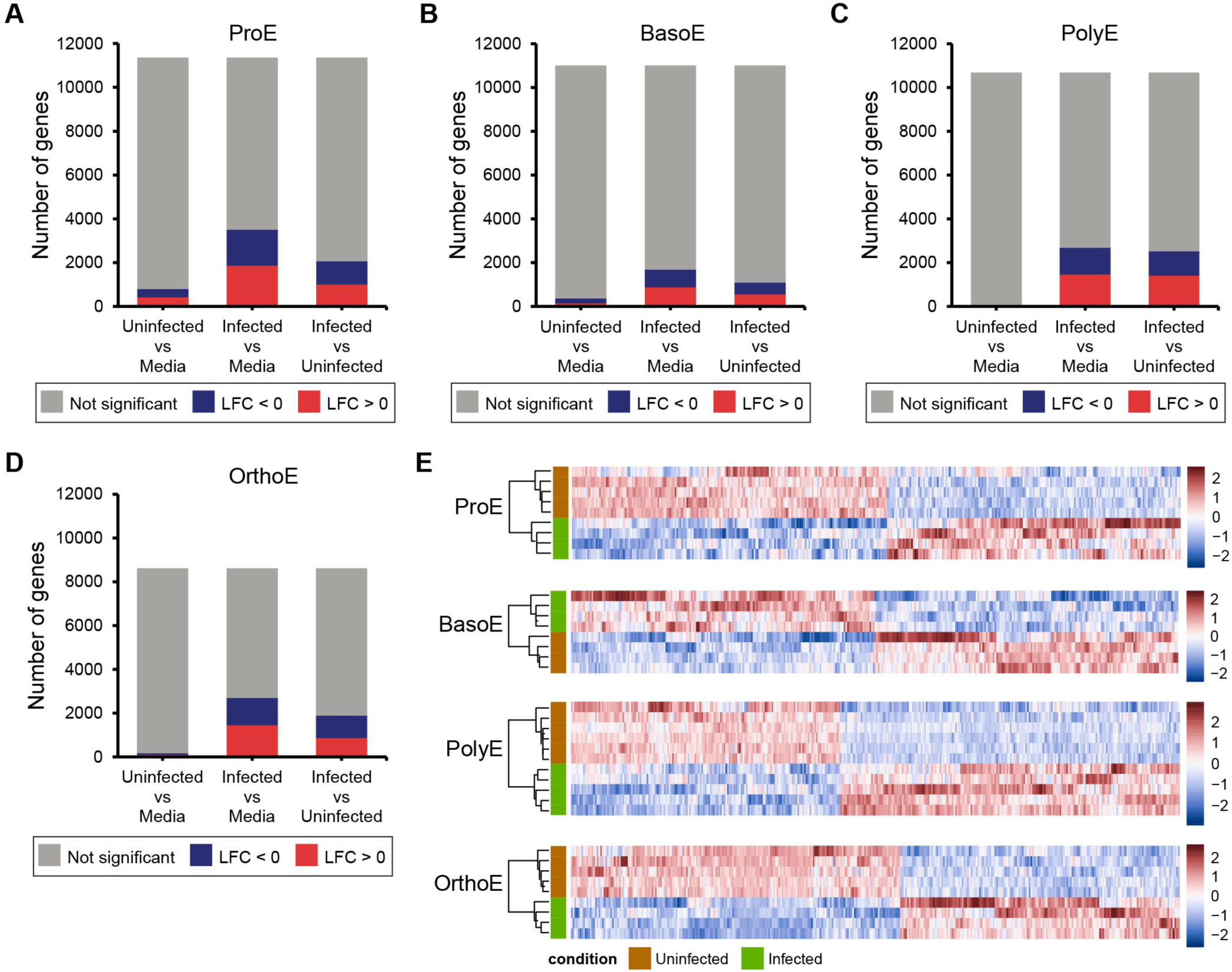
Differential gene expression detected in erythroblasts infected with *P. falciparum*. (A-D) Summaries of differential gene expression analysis for the pairwise comparison of populations by infection condition within each erythroblast cell type. Color indicates genes with log fold change > 0 (red), log fold change < 0 (blue), or non-significant fold change (gray). P-values were adjusted using the method of Benjamini and Hochberg and the threshold for significance set at 0.05. (E) Heatmaps showing z-score normalized expression of significant DEGs (infected vs uninfected) for each cell type.

In contrast, direct infection with *P. falciparum* produced host responses distinct from those found in uninfected erythroblasts or those exposed only to media (**Figure 3A-E**). Infection of ProE resulted in differential expression of 18% of detected genes compared to the uninfected population (**Figure 3A**). Only 10% of genes detected in BasoE were differentially expressed whereas we found differential expression of 24% of the PolyE transcriptome and 22% of the OrthoE transcriptome (**Figure 3B-D**). These findings suggest that the sensitivity of erythroblasts to parasite-induced transcriptional perturbation may vary by developmental stage. The proportion of upregulated and downregulated genes of the total DEGs was roughly equal regardless of the erythroblast stage (**Figure 3E**).

To dissect patterns of differential gene expression across cell types, we used set analysis to visualize the intersection of DEGs found by comparing the infected and uninfected transcriptomes of each erythroblast stage (**Figure 4**). The intersection of DEGs was considered separately for upregulated and downregulated genes. At least half the DEGs detected in each erythroblast stage, regardless of whether the DEGs had high or lower expression in infected erythroblasts, were unique to a single erythroblast stage. In contrast, only 27 DEGs were common to all four erythroblast stages. This result likely reflects the global changes in the composition of the erythroblast transcriptome over differentiation and suggests that the response to *P. falciparum* correlates with stage-specificity rather than a general, cell type-independent response to insult. The largest intersection in DEGs was observed between PolyE and OrthoE. 587 DEGs were unique to these two late erythroblast populations, whereas only 171 DEGs were common to ProE and BasoE. This could be due to fewer DEGs detected at the BasoE stage or a consequence of stage-specific differences related to the host response in early erythroblasts that are less pronounced at later stages.

**Figure 4:**
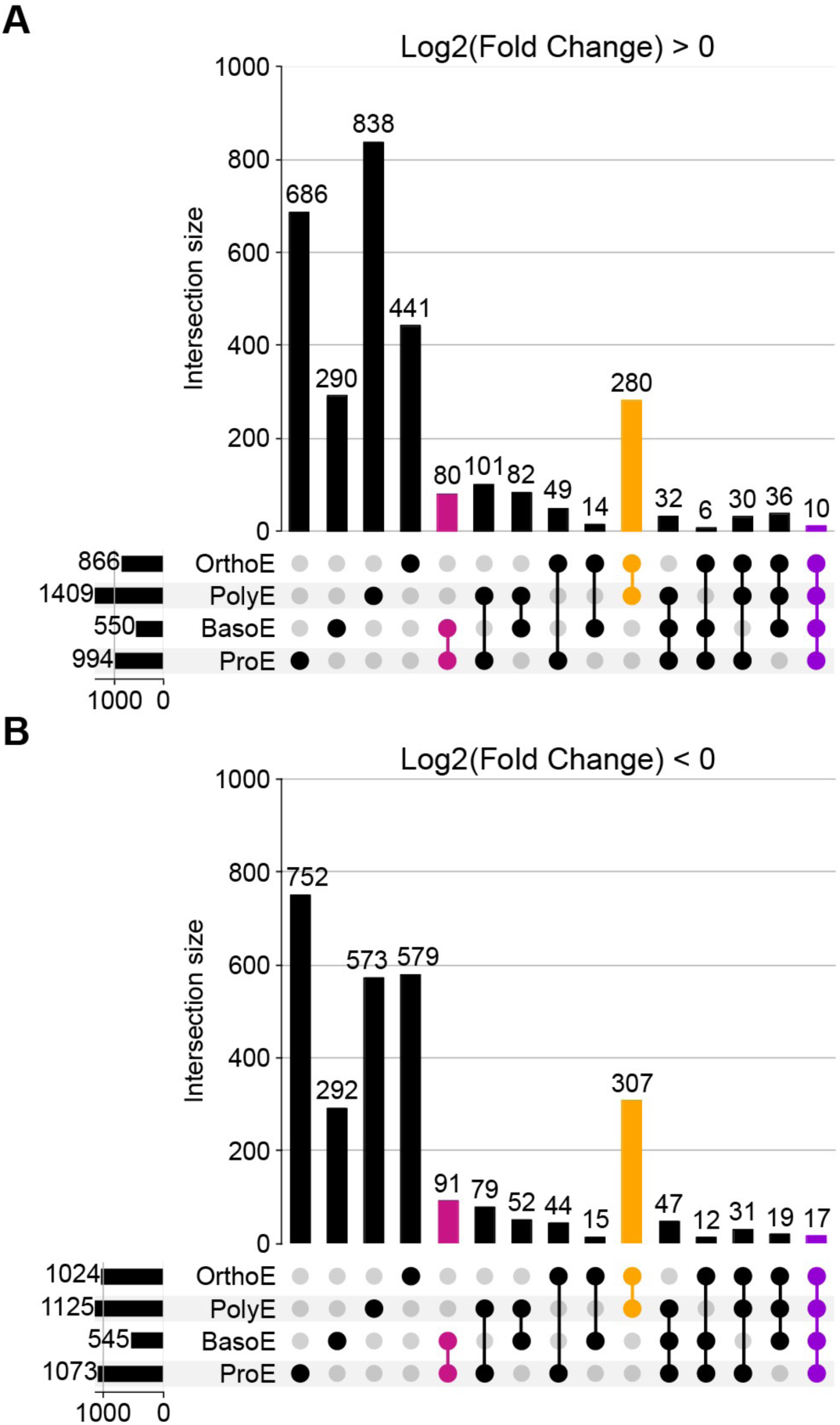
Erythroblasts have shared and distinct host responses to *P. falciparum* at different points along the erythroid trajectory. (A) Set intersections by cell type for DEGs that are more highly expressed in infected than uninfected erythroblasts (log fold change > 0). Color indicates set intersection for day 7 ProE and BasoE (magenta) and day 13 PolyE and OrthoE (orange), and all erythroblast populations (purple). (B) Set intersections for genes that have lower expression in infected than uninfected erythroblasts (log fold change < 0).

### Host responses in proerythroblasts and basophilic erythroblasts

Next, we used gene set analysis to characterize the host responses to *P. falciparum* infection in ProE and BasoE, which differ from PolyE and OrthoE in the composition and complexity of the transcriptome as well as hemoglobin content, a key factor in parasite nutrition. We applied enrichment analysis to identify cellular processes associated with differential gene expression in infected compared to uninfected erythroblasts (**Figure 5A, Supplementary Dataset 7**). EnrichR was used to determine if genes associated with processes in the MSigDB Hallmark 2020 gene set library were overrepresented among DEGs. Our analysis revealed that upregulated genes in the host response to infection of ProE and BasoE are significantly enriched for genes associated with E2F targets, G2M checkpoint, heme metabolism, and the mitotic spindle. None of the gene sets enriched among downregulated genes were common to ProE and BasoE.

**Figure 5:**
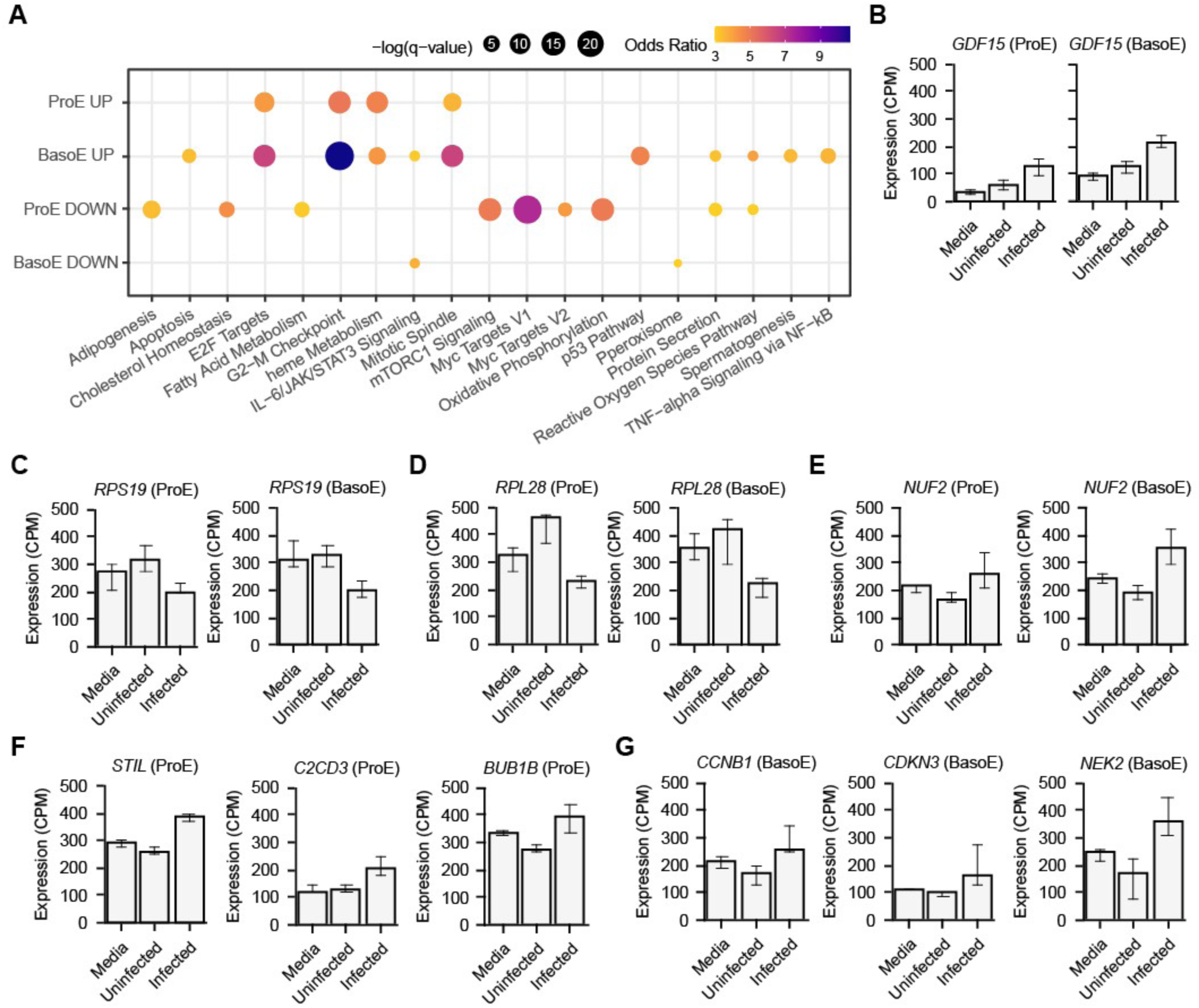
Host responses to infection with *P. falciparum* in early erythroblasts involve genes implicated in cell cycle regulation and dyserythropoiesis. (A) Enrichment analysis results using MSigDB Hallmark gene sets with DEGs (infected vs uninfected) detected in ProE and BasoE populations. Analysis of DEGs with log fold change > 0 are shown above the black line and analysis of DEGs with log fold change < 0 are shown below the black line. Size indicates significance. Color indicates the odds ratio from a Fisher’s exact test. (B) Expression of *GDF15* and (C) *NUF2* in ProE and BasoE populations. (D) Expression of cell-cycle related genes in ProE and (E) BasoE. (F-G) Expression of genes encoding ribosomal proteins in ProE and BasoE. Data are shown as median with 95% confidence intervals.

Notably, we also uncovered differential expression of genes implicated in the regulation of erythropoiesis in health and disease. *GDF15*, a member of the TGF-β superfamily and marker of ineffective erythropoiesis, was upregulated in infected ProE (2.1 fold, *p_adj_*= 8.6 ×10^-7^) and BasoE (1.8-fold, *p_adj_*=2.9×10^-6^) compared to uninfected neighboring cells (**Figure 5B**). Among downregulated genes in infected ProE and BasoE, we found several genes encoding ribosomal proteins. Decreased expression of transcripts for ribosomal proteins included *RPS19* and *RPL28*, genes implicated in pathogenesis of hereditary Diamond-Blackfan anemia (**Figure 5C, D**).

Perturbation of gene expression associated with the cell cycle and cell division was a common signature of infection. Upregulation of *NUF2*, a component of the kinetochore that is crucial for nuclear division, was common to both ProE and BasoE (2-fold, 6.5 ×10^-10^) (**Figure 5E**). Although the full set of genes associated with cell cycle and division function differed between ProE and BasoE, the host response of both cell types involved increased expression of regulators of cell division. In ProE, these included the mitotic regulator *STIL* (1.4-fold, *p_adj_*= 7.4 ×10^-11^), centriole elongator regulator *C2CD3* (1.6-fold, *p_adj_*= 1.2 ×10^-7^), and spindle-assembly regulator *BUB1B* (1.4-fold, *p_adj_=* 1.2 ×10^-5^) (**Figure 5F**). In BasoE, we found upregulation of cyclin B1 *CCNB1* (1.7-fold, *p_adj_*= 3.9 ×10^-5^), cyclin dependent kinase inhibitor *CDKN3* (2-fold, *p_adj_=* 4.0 ×10^-5^), and mitotic regulator *NEK2* (2.4-fold, *p_adj_*= 1.6 ×10^-7^) (**Figure 5G**). Together, the observed host responses indicate that *P. falciparum* infection at the start of terminal differentiation induces host responses that are common to the pathogenesis of dyserythropoiesis in anemia of inflammation as well as hereditary disorders of hematopoiesis.

### Host responses in polychromatic and orthochromatic erythroblasts

We hypothesized that PolyE and OrthoE would respond differently to infection than early erythroblasts, reasoning that terminal differentiation would affect host defenses and the capacity to support parasite development. Enrichment analysis uncovered gene sets that were overrepresented in the host response of PolyE and OrthoE to direct infection with *P. falciparum*, compared to uninfected neighboring cells (**Figure 6A, Supplementary Dataset 8**). The upregulated DEGs in infected PolyE were enriched for genes belonging to the apoptosis and protein secretion gene sets whereas downregulated DEGs were enriched for genes related to E2F targets, G2-M checkpoint, and the mitotic spindle. In contrast, members of the E2F targets, G2-M checkpoint, and Myc targets V1 and V2 were overrepresented in DEGs upregulated in infected OrthoE. Both populations of late erythroblasts upregulated genes associated with mTORC1 signaling, the p53 pathway, TNF-a signaling via the NFkB pathway, the unfolded protein response, and the UV response. We also found downregulation of genes involved in heme metabolism and management of reactive oxygen species. Both processes are crucial for red cell development.

**Figure 6:**
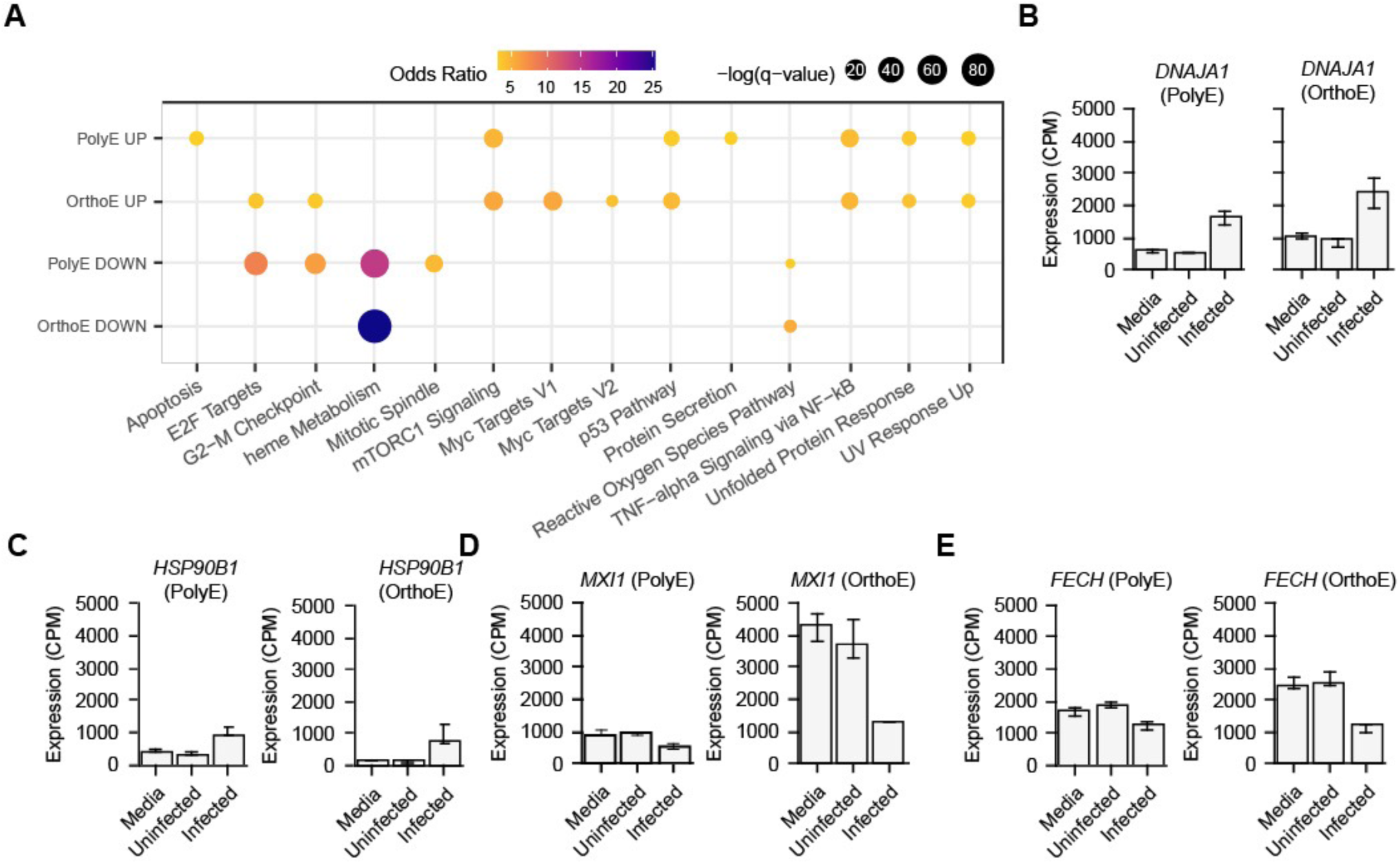
Host responses to infection with *P. falciparum* in late erythroblasts. (A) Enrichment analysis results using MSigDB Hallmark gene sets with DEGs (infected vs uninfected) detected in PolyE and OrthoE populations. Analysis of DEGs with log fold change > 0 are shown above the black line and analysis of DEGs with log fold change < 0 are shown below the black line. Size indicates significance. Color indicates the odds ratio from a Fisher’s exact test. (B,C) Expression of stress-related genes in PolyE and OrthoE populations. (D) Expression of *MXI1*, an enucleation factor, in PolyE and OrthoE. (E) Expression of *FECH*, encoding protoporphyrin ferrochetalase, in PolyE and OrthoE. Data are shown as median with 95% confidence intervals.

Among the shared responses between PolyE and OrthoE were genes involved in proteotoxic stress and several genes that are essential for the final stages of erythroid differentiation. We found increased expression of *DNAJA1*, a member of the Hsp40 family of protein chaperones, in infected PolyE (2.8-fold, *p_adj_*= 4.5 ×10^-18^) and OrthoE (1.6-fold, *p_adj_*= 6.9 ×10^-4^) (**Figure 6B**). Expression of the *HSP90B1* gene, which encodes a chaperone whose client proteins include certain integrins and toll-like receptors, was also increased in infected PolyE (2.7-fold, *p_adj_*= 8.7 ×10^-21^) and OrthoE (3.6-fold, *p_adj_*= 9.8 ×10^-8^) (**Figure 6C**). Perturbation of genes involved in healthy erythroid development was also evident from our analysis. *MXI1*, a critical factor for enucleation, was downregulated in PolyE (1.7-fold, *p_adj_*= 3.6 ×10^-11^) and OrthoE (4.9-fold, *p_adj_*= 4.2 2×10^-50^) (**Figure 6D**). Knockdown of *MXI1* expression in murine erythroblasts reduces enucleation and results in decreased expression of other genes involved in erythroid development such as *FECH1*, the murine homolog of the human *FECH* gene.^38^ Expression of *FECH*, which encodes a key enzyme in heme biosynthesis, was indeed decreased 1.6-fold in infected PolyE (1.6-fold, *p_adj_*= 1.6 ×10^-10^) and greater than 3-fold in infected OrthoE (3.7-fold, *p_adj_*= 8.6×10^-28^) (**Figure 6E**). Deficient activity of the ferrochetalase encoded by *FECH* is implicated in hereditary erythropoietic protoporphyria^39^, highlighting the detrimental effect of perturbed heme metabolism on the host.

### Erythroblasts at all stages upregulate HMOX1 in response to infection with P. falciparum

Although many changes in the host transcriptome were dependent on erythroblast stage, we found that direct infection induced changes to genes involved in mitigating oxidative stress in a stage-independent manner. *HMOX1*, which encodes heme oxygenase 1, was upregulated in each population of infected erythroblasts **(Figure 7**). We found a 3.6-fold increase in expression in infected ProE (*p_adj_*=3.9×10^-9^) and a 5.9-fold increase in infected BasoE (*p_adj_*=4.0×10^-4^). Expression of *HMOX1* was increased 19-fold in infected PolyE (*p_adj_*=2.3×10^-29^) and 6.5-fold in infected OrthoE (*p_adj_*= 1.6 ×10^-27^). These data show that *HMOX1* expression is involved in the erythroblast response to *P. falciparum* at all stages of terminal differentiation, adding to previous evidence that *HMOX1* is a critical host factor in malaria infection.^40–42^

**Figure 7:**
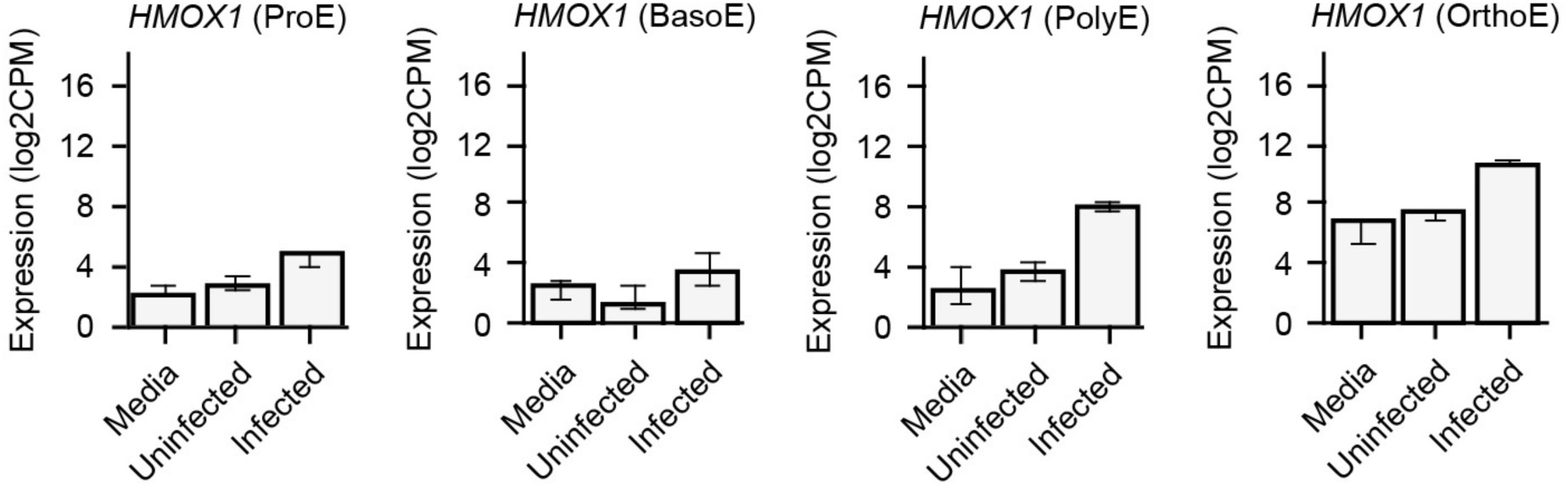
Upregulation of *HMOX1* in erythroblasts directly infected with *P. falciparum.* Expression of *HMOX1*, encoding heme oxygenase-1, in each erythroblast population. Data are shown as median with 95% confidence intervals.

### Genes encoding parasite effectors that are exported into the host cytoplasm are transcribed in erythroblast infection

*P. falciparum* extensively modifies the host red blood cell and hepatocytes by exporting effectors into the host cytoplasm,^43–45^ and related parasites are known to modulate host gene expression through effectors that traffic to the host cell nucleus.^46, 47^ We hypothesized that the same might be true in *P. falciparum* during infection of nucleated erythroblasts, so we queried our RNA-seq dataset for transcription of genes encoding the predicted intraerythrocytic “exportome” of *P. falciparum*.^36, 48^ We found that of 245 transcripts of exported genes expressed at 8-or 16-hours post-invasion in RBCs, 185 were expressed in parasites in at least one of the four erythroblast stages (**Supplementary Table S2**). Among those detected were genes encoding well-studied examples of exported effectors, including ring-infected erythrocyte surface antigen (PF3D7_0102200), knob-associated histidine-rich protein (PF3D7_0202000), and members of the FIKK kinase family. These data provide a foundation for further study of effector export during infection of erythroblasts by *P. falciparum*.

## Discussion

Although dyserythropoiesis has long been recognized as a contributing cause in severe malarial anemia, little is understood about the host-parasite interactions that lead to perturbed erythroid development. In this study, we employed *ex-vivo* erythropoiesis of primary human bone marrow CD34^+^ HSPCs to study transcriptional host responses to infection by *P. falciparum* asexual parasites at specific stages of erythroid development. Using a small panel of erythroid surface markers enabled us to sort and sequence the transcriptome of infected and uninfected erythroblast populations at four distinct stages of terminal differentiation.

We found few DEGs from the comparison of uninfected cells to cells cultured in media without stimulus or SRS. This result suggests that ∼20-22 hours post-invasion of erythroblasts, transcriptional changes due to signaling from infected cells to neighboring uninfected cells or from exposure to releasate from ruptured schizonts were negligible. This type of paracrine signaling may be more relevant at earlier timepoints, and it is possible that activity by the parasite silences signaling from infected cells to uninfected neighbors, as has been found in a related member of the apicomplexan parasite family, *Toxoplasma gondii*.^47^ Although Lamikanra *et al.* observed differential gene expression between hemozoin-treated erythroblasts compared to controls, the concentration of hemozoin used was much greater than that generated by the MOI used in this study.^34^ Further work is warranted to better understand the implications for erythroid development from the presence of hemozoin and other inflammatory byproducts of schizont rupture in the bone marrow.

Many of the DEGs we found in infected erythroblasts compared with uninfected cells are implicated in hereditary anemias whose pathogenesis involves disordered erythropoiesis. Expression of GDF15, a regulator of stress erythropoiesis, could have systemic effects on regulation of iron metabolism as is found in patients with *β*-thalassemia.^49^ We also found altered expression of transcripts encoding ribosomal proteins, a hallmark of Diamond-Blackfan Anemia (DBA). DBA is an inherited condition caused by mutations in genes that encode ribosomal proteins and characterized by macrocytic anemia, physical abnormalities, and a lack of erythroid precursors in bone marrow with otherwise normal cellularity.^50^ Decreased expression of *RPS19*, a common etiology of DBA, has been shown to reduce *in vitro* proliferation of erythroid progenitor cells through activation of the p53 pathway and cell cycle arrest.^51, 52^ Recent work on regulation of hematopoiesis has underscored that not only does expression of individual ribosomal components affect cell development, but the abundance of ribosomes can determine cell fate in stem cell precursors.^53^ Disruption of the sensitive balance between proliferation of precursor cells and erythroid commitment and differentiation, further evidenced by differential expression of genes associated with the cell division, could have a similar detrimental effect on production of healthy erythrocytes in the malaria-infected bone marrow.

Our study also indicates that parasite infection induces changes in expression of genes related to protein-folding pathways and factors that mitigate proteotoxic stress. In infected PolyE and OrthoE, we found increased transcription of genes encoding protein chaperones like *DNAJA1* and *HSP90B1*. Protein chaperones function at several critical steps in erythropoiesis from shuttling cyclins into the nucleus and cooperating with cyclin-dependent kinases to regulate cell division to facilitating globin folding and assembly and preventing premature apoptosis in response to the unusual demands of remodeling the proteome in erythrocytic maturation.^54^ As such, dysregulation of members of the HSP70 family and other molecular chaperones is often implicated in disordered erythropoiesis. In DBA, mutations in *RPL5* or *RPL11* result in decreased levels of HSP70 leading to excessive degradation of GATA1 and defects in proliferation, differentiation, and regulation of apoptosis.^55^ Overexpression of HSP70 rescues DBA progenitors by increasing GATA1 levels and reducing free heme by restoring normal hemoglobin synthesis.^56^ HSP90 is also connected to protection from cellular stress as a chaperone of FANCA, which participates in the DNA damage response. Mutation of FANC genes is the underlying etiology in Fanconi anemia, a rare disorder characterized by genomic instability, bone marrow hypocellularity, and subsequent pancytopenia of the three blood cell lineages.^57^ The increased transcription of genes encoding heat shock proteins suggest that *Plasmodium* infection of late erythroblasts may produce genotoxic or proteotoxic effects that result in dyserythropoiesis. Although further experiments are needed to evaluate the extent of damage to the cell during infection, increased transcription suggests a cell intrinsic host response aimed at protecting developing erythroblasts from detrimental effects of infection.

In the peripheral blood, *P. falciparum* extensively modifies infected red cells by exporting proteins from the parasitophorous vacuole (PV) where the parasite resides after invasion and throughout its development.^45^ *Plasmodium* has also been shown to export proteins into the hepatocyte cytoplasm during the liver stage of infection, some of which reach the host cell nucleus, although the full exportome and activity of effectors remain largely unknown.^43, 44^ It is possible that *Plasmodium* effectors are similarly involved in mediating host cell responses to infection and transforming the erythroblast into a hospitable environment for gametocytogenesis. Given that we observed differential expression of genes that regulate enucleation and heme metabolism, it is tempting to speculate that parasite activity is involved in physical and metabolic modification of the host cell.

In this study, we demonstrated that erythroblasts have specific transcriptional responses to infection with *P. falciparum* at distinct stages of erythropoiesis. Moreover, direct infection with *P. falciparum* induces differential expression of genes related to cell cycle regulation and key erythroid developmental processes in addition to a wide range of stress responses. The results presented here provide a foundation for further investigation of the impact of malaria infection on erythropoiesis and the pathogenesis of malaria anemia. A more complex model of the hematopoietic niche that better mimics the microenvironment of erythroblastic islands and a single cell approach to transcriptomics could reveal the broader impact of parasite infection on the development of blood cell lineages in the hematopoietic niche.^20^ Genetic manipulation of *ex-vivo* differentiated erythroblasts could be used to interrogate host factors identified in this study or the effect of naturally occurring genotypes in regions of malaria endemicity, like sickle hemoglobin, on erythroblast responses to *P. falciparum*. Further elaboration of the interactions between host and parasite in the bone marrow may yield strategies for mitigating harm to host cell development and limiting the bone marrow as a reservoir for growth and development of the parasite.

## Acknowledgements

We thank David Schneider, Anupama Narla, John Boothroyd, and members of the Egan, Boothroyd, and Yeh labs for helpful discussions. This work was supported in part by NIH T32GM007276 (T.P.F)., NIH DP2HL13718601 (E.S.E.), and the Stanford Maternal Child Health Research Institute. Elizabeth S. Egan, M.D., Ph.D. holds an Investigators in the Pathogenesis of Infectious Disease Award from the Burroughs Wellcome Fund, and is a Chan Zuckerberg Biohub-San Francisco Investigator and a Tashia and John Morgridge Endowed Faculty Scholar of the Stanford Maternal and Child Health Research Institute. This work used the Genome Sequencing Service Center by Stanford Center for Genomics and Personalized Medicine Sequencing Center, supported by the grant award NIH S10OD020141. Some of the computing for this project was performed on the Sherlock and SCG clusters. We would like to thank Stanford University and the Stanford Research Computing Center for providing computational resources and support that contributed to these research results. Parts of Figure 1B and Figure 2A were created with Biorender.com.

## Authorship Contributions

Conceptualization, T.P.F. and E.S.E.; Performed experiments, T.P.F., Y.R., and E.S.E.; Data analysis and visualization, T.P.F; Resources, E.S.E.; Writing-initial draft, T.P.F.; Writing-final draft, T.P.F and E.S.E.; Supervision, E.S.E.; Funding acquisition, E.S.E.

## Disclosure of Conflicts of Interest

The authors have no conflicts of interest related to the work presented in this manuscript.

## Notes

### Competing Interest Statement

The authors have declared no competing interest.

### Summary of Updates

Revised figures/tables after a duplicated row was noted in differential expression table used for analyses. Outcomes not impacted.

