## Supplementary material for "*Plasmodium falciparum* infection of human erythroblasts induces transcriptional changes associated with dyserythropoiesis"

**This PDF file includes:**

Supplemental Methods

Supplemental Figures: Figures S1 and S2

Supplemental Tables: Tables S1 and S2

**Other supplementary material for this manuscript includes the following:**

Datasets S1 to S8 are submitted as separate Excel files.

**Supplemental Methods**

**Primary human cell culture and differentiation**

CD34+ human bone marrow cells (STEMCELL Technologies) were maintained in static cultures at 37ºC in 5% CO2. Cells were grown in IMDM media (Iscove's Modified Dulbecco's Medium with stable glutamine, Sigma) with the addition 330 ug/mL holo-transferrin (BBI Solutions), 10 ug/mL heparin (Affymetrix), 10 ug/mL recombinant human insulin (Stem Cell Tech), 0.5% v/v of 10 kU/mL penicillin/streptomycin solution (Gibco) and 0.5% human plasma (Octaplas, Octapharma). Media was supplemented with the following cytokines according to the stage of culture: 10-6 M hydrocortisone (Sigma), 100 ng/mL Stem Cell Factor (SCF, R&D Systems), 5 ng/mL IL-3 (R&D Systems), 3 IU/mL erythropoietin (EPO, Amgen).

Cultures were initiated with 1 x $10^6$ $CD34^+$ cells. On day 0, cells were thawed, washed twice, and seeded at 1.3 x $10^4$ cells/mL in media supplemented with hydrocortisone, IL-3, SCF, and EPO. On day 4, cells were washed once and diluted 1:5 in fresh media. On day 7, cells were washed once and plated at 4 x $10^5$ cells/mL in fresh media supplemented with SCF and EPO. Cultures were diluted to a concentration of 5 x $10^5$ cells/mL in fresh media on day 9 of differentiation. On day 11 of differentiation, cells were washed once and plated in fresh media with EPO at 1 x $10^6$/mL. Cells were maintained in media with EPO until the end of culture.

**Infection of primary human erythroblasts**

Infections were carried out in a modified version of the primary culture medium described above. Infection media lacked heparin, an inhibitor of parasite invasion, and included 50 mg/L hypoxanthine to support parasite growth. Instead of human plasma, media was supplemented with 10% Albumax. Cytokines were included according to the day of differentiation of the erythroblast culture as described above.

On the day of infection, erythroblasts were removed from the parent culture and washed twice. Erythroblasts were resuspended in infection media with cytokines and 2 x $10^6$ erythroblasts were plated per well in six-well plates on day 7 and 4.5 x $10^6$ erythroblasts were plated per dish in 6 cm dishes on day 13. Six replicate wells or dishes were prepared per experimental condition and treated as biological replicates for downstream analyses. Tightly synchronized schizont parasites of strain D10-pfPHG were purified by magnetic column (Miltenyi biotec) and resuspended in infection media with cytokines. Schizonts were mixed with day 7 erythroblasts at a ratio of 5:1 and with day 13 erythroblasts at a ratio of 3:1. For the syringe-ruptured schizont (SRS) condition, schizonts were passed through a 1.2 µm filter and incubated at room temperature for 15 minutes to ensure no viable merozoites were present before mixing with erythroblasts. Media was added to wells for unstimulated erythroblasts to an equivalent volume of the SRS and live parasite wells. Additional schizonts from the purification were put back in erythrocyte culture to monitor egress. At ~5h post-egress, half the volume was removed from each well and replaced with fresh media.

**FACS and analysis of erythroblast populations**

Cells were prepared for analysis by washing twice in 1X PBS with 0.3% BSA. For experiments in which erythroblasts were staged, the following staining cocktail was used: 1:30 CD235a-APC eFluor780 clone HIR2 (ThermoFisher, 47-9987-42), 1:100 CD233-PE clone BRIC6 (IGBRL, 9439), CD49d-APC clone MZ18-24A9 (Miltenyi, 130-093-279), 1 uM Calcein Violet 450 AM (ThermoFisher, 65-0854-39). Staining was performed on ice for 30 minutes. After staining, cells were washed twice and resuspended in 1X PBS with 0.3% BSA. Data collection and cell sorting was performed on a FACSAriaII (BD Biosciences) in the Stanford Shared FACS Facility. For microscopy, cells were stained only with 1 ug/mL Hoechst 33342 (ThermoFisher) and sorted into FBS for visualization on a Keyence BZ-X700 fluorescence microscope with the DAPI (OP-87762) and GFP (OP-87763) filter sets. Cells used for RNA-extraction were sorted directly into Buffer RLT (Qiagen) with beta-mercaptoethanol according to the manufacturer's instructions.

**Data processing and read alignment**

Custom scripts in Bash, Python, and R were used for analysis. Sequencing data was demultiplexed using bcl2fastq (Illumina) and assessed for quality using FastQC (https://www.bioinformatics.babraham.ac.uk/projects/fastqc/). Replicates that failed QC were removed from analyses. Adaptor trimming was performed using TrimGalore (https://www.bioinformatics.babraham.ac.uk/projects/trim_galore/). BBTools suite (BBMap – Bushnell, B. - https://www.sourceforge.net/projects/bbmap/) was used to remove human and parasite rRNA reads. Reads were aligned to a concatenation of the human (GRCh38) and parasite (ASM276v2) genomes using STAR with default settings. Gene counts were generated by running the -quantMode flag in STAR.

**Principal component analysis (PCA)**

Parasite genes were not included in the table of counts used for PCA. For PCA, counts were transformed using the vst function in DESeq2 and analysis was carried out using the prcomp function in the R stats package with default settings.

**Differential Gene Expression**

DESeq2 was used to conduct differential gene expression analysis on the host genes between erythroblast populations by erythroblast stage. P-value adjustment was performed by the DESeq2 implementation of independent hypothesis weighting with an alpha of 0.05 for controlling the false discovery rate. Heatmap visualization of differentially expressed genes was performed using the pheatmap package on counts transformed by the vst function and scaled per gene using the z-score. The samples were clustered by Ward's method using the pheatmap flag clustering_method set to ward.D2. Set analysis was conducted and visualized using the upsetplot Python package (https://upsetplot.readthedocs.io/en/stable/). Expression of individual genes was plotted as counts per million (CPM) or log2(CPM).

**Enrichment analysis and visualization**

Analysis for gene set enrichment was performed using the Independent Enrichment Analysis online tool (https://appyters.maayanlab.cloud/Independent_Enrichment_Analysis/) with the MSigDB Hallmark 2020 gene sets. The background set was composed of all protein-coding genes. An adjusted p-value cutoff was set at 0.1 and a minimum odds ratio of 3 was used to filter significant results.

**Analysis of parasite gene expression**

Parasite gene counts were normalized by DESeq2's median of ratios method and averaged across replicates for each erythroblast stage. The reference "exportome" was sourced from Jonsdottir *et al.*[48] and filtered to exclude genes which are not transcribed at the 8-16h timepoints in the blood stage transcriptome published by Otto *et al*.[36]

### Supplementary Figures

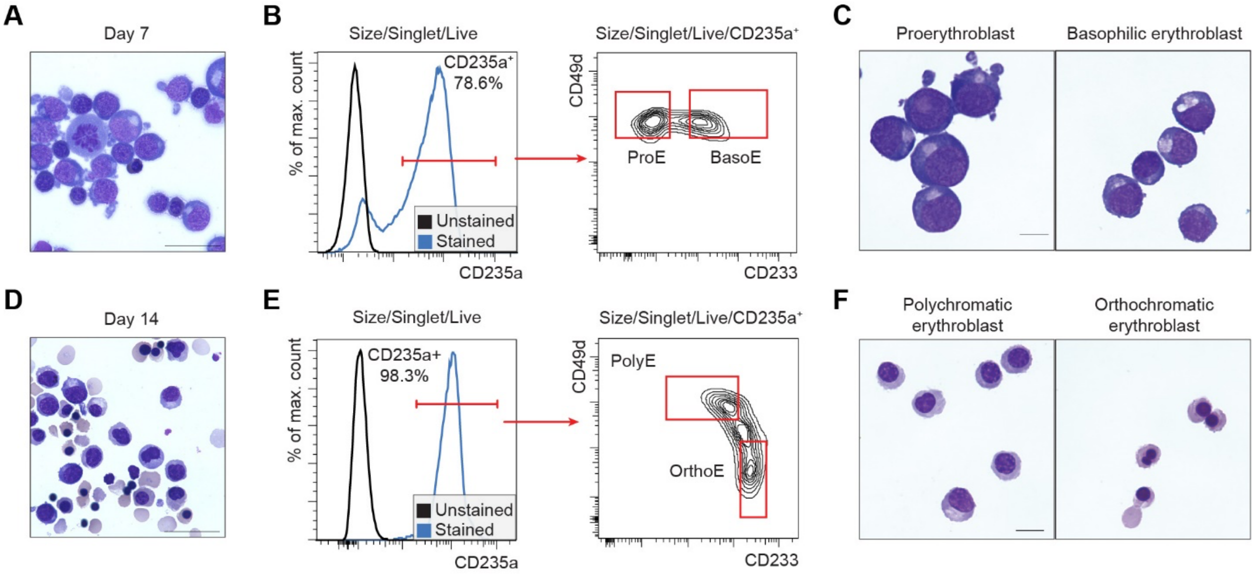


**Figure S1: Staging and isolation of erythroblast populations by Fluorescence Activated Cell Sorting *(FACS*).** (A) Representative cytospin of primary human erythroblasts on day 7 of ex vivo growth. Scale bar represents 50µm. (B) Flow cytometry gating strategy for identifying proerythroblasts (ProE) and basophilic erythroblasts (BasoE) based on expression of CD235a, CD233, and CD49d. (C) Representative cytospins of day 7 erythroblast populations after FACS. Scale bar represents 10µm. (D) Representative cytospin of primary human erythroblasts on day 14 of ex vivo growth. Scale bar represents 50µm. (E) Flow cytometry gating strategy for identifying polychromatic erythroblasts (PolyE) and orthochromatic erythroblasts (OrthoE) based on expression of CD235a, CD233, and CD49d. (F) Representative cytospins of day 14 erythroblast populations after FACS. Scale bar represents 10µm.

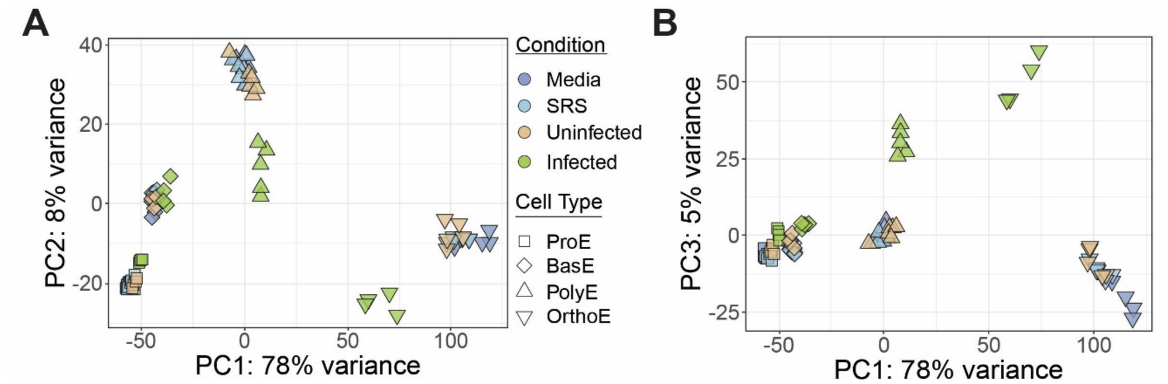


**Figure S2: Variance between media-only, syringe-lysed schizonts (SRS), and uninfected (GFP$^{-}$) conditions is minimal.** (A) Principal component analysis of RNA-seq dataset. Marker shape indicates erythroblast population. Color represents infection condition. Two principal components (PCs) are plotted, PC1 and PC2. (B) As in (A) for PC1 and PC3.

SUPPLEMENTARY TABLE 1: Summary of total, *H. sapiens* , and *P. falciparum* gene counts

| Library | Total gene counts | *H. sapiens* gene counts | *H. sapiens* gene counts (% of total) | *P. falciparum* gene counts | *P. falciparum* gene counts (% of total) |
|---|---|---|---|---|---|
| ProE.Media.11 | 9876559 | 9872333 | 99.96 | 4226 | 0.04 |
| ProE.Media.12 | 9183465 | 9180166 | 99.96 | 3299 | 0.04 |
| ProE.Media.21 | 8268678 | 8265340 | 99.96 | 3338 | 0.04 |
| ProE.Media.22 | 10522131 | 10518375 | 99.96 | 3756 | 0.04 |
| ProE.Media.31 | 7958466 | 7954837 | 99.95 | 3629 | 0.05 |
| ProE.Media.32 | 8409436 | 8406684 | 99.97 | 2752 | 0.03 |
| ProE.SRS.11 | 7035612 | 7032891 | 99.96 | 2721 | 0.04 |
| ProE.SRS.12 | 7682074 | 7679349 | 99.96 | 2725 | 0.04 |
| ProE.SRS.21 | 7733241 | 7729990 | 99.96 | 3251 | 0.04 |
| ProE.SRS.22 | 7461290 | 7458636 | 99.96 | 2654 | 0.04 |
| ProE.SRS.31 | 9237393 | 9232161 | 99.94 | 5232 | 0.06 |
| ProE.SRS.32 | 11495784 | 11492700 | 99.97 | 3084 | 0.03 |
| ProE.Uninfected.12 | 9486756 | 9482304 | 99.95 | 4452 | 0.05 |
| ProE.Uninfected.21 | 8889118 | 8883199 | 99.93 | 5919 | 0.07 |
| ProE.Uninfected.22 | 9014061 | 9010230 | 99.96 | 3831 | 0.04 |
| ProE.Uninfected.31 | 8016362 | 8010804 | 99.93 | 5558 | 0.07 |
| ProE.Uninfected.32 | 9667211 | 9662529 | 99.95 | 4682 | 0.05 |
| ProE.Infected.21 | 9844754 | 9580848 | 97.32 | 263906 | 2.68 |
| ProE.Infected.22 | 8952267 | 8699997 | 97.18 | 252266 | 2.82 |
| ProE.Infected.31 | 13400200 | 12990191 | 96.94 | 410009 | 3.06 |
| ProE.Infected.32 | 7705835 | 7418256 | 96.27 | 287579 | 3.73 |
| BasoE.Media.11 | 7110193 | 7107195 | 99.96 | 2998 | 0.04 |
| BasoE.Media.12 | 8306751 | 8302719 | 99.95 | 4032 | 0.05 |
| BasoE.Media.21 | 8395734 | 8391617 | 99.95 | 4117 | 0.05 |
| BasoE.Media.22 | 8518715 | 8511797 | 99.92 | 6918 | 0.08 |
| BasoE.Media.31 | 8093837 | 8089588 | 99.95 | 4249 | 0.05 |
| BasoE.Media.32 | 7441416 | 7438212 | 99.96 | 3204 | 0.04 |
| BasoE.SRS.11 | 7753775 | 7750087 | 99.95 | 3688 | 0.05 |
| BasoE.SRS.12 | 7362136 | 7359089 | 99.96 | 3047 | 0.04 |
| BasoE.SRS.21 | 7863215 | 7858198 | 99.94 | 5017 | 0.06 |
| BasoE.SRS.22 | 9292865 | 9290135 | 99.97 | 2730 | 0.03 |
| BasoE.SRS.31 | 9496652 | 9487657 | 99.91 | 8995 | 0.09 |
| BasoE.SRS.32 | 8243895 | 8239438 | 99.95 | 4457 | 0.05 |
| BasoE.Uninfected.21 | 11097709 | 11091132 | 99.94 | 6577 | 0.06 |
| BasoE.Uninfected.22 | 8397371 | 8391884 | 99.93 | 5487 | 0.07 |
| BasoE.Uninfected.31 | 9085282 | 9080366 | 99.95 | 4916 | 0.05 |
| BasoE.Uninfected.32 | 9004602 | 8996643 | 99.91 | 7959 | 0.09 |
| BasoE.Infected.12 | 9315109 | 8805290 | 94.53 | 509819 | 5.47 |
| BasoE.Infected.22 | 8462830 | 7999104 | 94.52 | 463726 | 5.48 |
| BasoE.Infected.31 | 9399368 | 8947677 | 95.19 | 451691 | 4.81 |
| BasoE.Infected.32 | 11778349 | 11096474 | 94.21 | 681875 | 5.79 |
| PolyE.Media.11 | 10122971 | 10118456 | 99.96 | 4515 | 0.04 |
| PolyE.Media.12 | 8373718 | 8370081 | 99.96 | 3637 | 0.04 |
| PolyE.Media.21 | 8204941 | 8202202 | 99.97 | 2739 | 0.03 |
| PolyE.Media.22 | 10856134 | 10852372 | 99.97 | 3762 | 0.03 |
| PolyE.Media.31 | 14982652 | 14978389 | 99.97 | 4263 | 0.03 |
| PolyE.SRS.11 | 9528939 | 9524957 | 99.96 | 3982 | 0.04 |
| PolyE.SRS.12 | 9720557 | 9715678 | 99.95 | 4879 | 0.05 |
| PolyE.SRS.21 | 9253393 | 9249777 | 99.96 | 3616 | 0.04 |
| PolyE.SRS.22 | 7351203 | 7348424 | 99.96 | 2779 | 0.04 |
| PolyE.SRS.31 | 8013726 | 8010521 | 99.96 | 3205 | 0.04 |
| PolyE.SRS.32 | 11346144 | 11341948 | 99.96 | 4196 | 0.04 |
| PolyE.Uninfected.11 | 8820693 | 8814943 | 99.93 | 5750 | 0.07 |
| PolyE.Uninfected.12 | 10203527 | 10196466 | 99.93 | 7061 | 0.07 |
| PolyE.Uninfected.21 | 11519894 | 11500595 | 99.83 | 19299 | 0.17 |
| PolyE.Uninfected.22 | 10997346 | 10988389 | 99.92 | 8957 | 0.08 |
| PolyE.Uninfected.31 | 9609616 | 9603506 | 99.94 | 6110 | 0.06 |
| PolyE.Uninfected.32 | 9895331 | 9891010 | 99.96 | 4321 | 0.04 |
| PolyE.Infected.11 | 9145366 | 8281710 | 90.56 | 863656 | 9.44 |
| PolyE.Infected.12 | 8684484 | 8120134 | 93.5 | 564347 | 6.5 |
| PolyE.Infected.21 | 8894665 | 8382929 | 94.25 | 511736 | 5.75 |
| PolyE.Infected.22 | 9853765 | 8594081 | 87.22 | 1259684 | 12.78 |
| PolyE.Infected.31 | 11018562 | 8618244 | 78.22 | 2400317 | 21.78 |
| OrthoE.Media.11 | 9007715 | 9003854 | 99.96 | 3861 | 0.04 |
| OrthoE.Media.12 | 11230384 | 11226187 | 99.96 | 4197 | 0.04 |
| OrthoE.Media.21 | 10288349 | 10283361 | 99.95 | 4988 | 0.05 |
| OrthoE.Media.22 | 10265363 | 10261647 | 99.96 | 3716 | 0.04 |
| OrthoE.Media.32 | 8548602 | 8545594 | 99.96 | 3008 | 0.04 |
| OrthoE.SRS.11 | 13097323 | 13090340 | 99.95 | 6983 | 0.05 |
| OrthoE.SRS.12 | 12397321 | 12391683 | 99.95 | 5638 | 0.05 |

| | | | | | |
|---|---|---|---|---|---|
| OrthoE.SRS.21 | 7827098 | 7822244 | 99.94 | 4854 | 0.06 |
| OrthoE.SRS.22 | 9216537 | 9211779 | 99.95 | 4758 | 0.05 |
| OrthoE.SRS.31 | 8241174 | 8236430 | 99.94 | 4744 | 0.06 |
| OrthoE.SRS.32 | 8404283 | 8398753 | 99.93 | 5530 | 0.07 |
| OrthoE.Uninfected.11 | 7777553 | 7744190 | 99.57 | 33363 | 0.43 |
| OrthoE.Uninfected.12 | 9624433 | 9586510 | 99.61 | 37923 | 0.39 |
| OrthoE.Uninfected.21 | 7945723 | 7916992 | 99.64 | 28731 | 0.36 |
| OrthoE.Uninfected.22 | 7838152 | 7812489 | 99.67 | 25663 | 0.33 |
| OrthoE.Uninfected.32 | 7819867 | 7795968 | 99.69 | 23899 | 0.31 |
| OrthoE.Infected.11 | 18536726 | 7970335 | 43 | 10566388 | 57 |
| OrthoE.Infected.12 | 21557918 | 9931817 | 46.07 | 11626099 | 53.93 |
| OrthoE.Infected.21 | 19811460 | 9617828 | 48.55 | 10193630 | 51.45 |
| OrthoE.Infected.22 | 19624274 | 8617038 | 43.91 | 11007235 | 56.09 |

SUPPLEMENTARY TABLE 2: Transcriptional expression of the intraerythrocytic parasite exportome during erythroblast infection

| Gene ID | Description | PEXEL.motif | Localisation in infected erythrocytes |
|---|---|---|---|
| PF3D7_0102200 | ring-infected erythrocyte surface antigen | RNLYGE | iRBC periphery |
| PF3D7_0102600 | serine/threonine protein kinase, FIKK family (FIKK1) | RYLAE | |
| PF3D7_0104200 | StAR-related lipid transfer protein | RILKE | PV, iRBC |
| PF3D7_0112900 | Plasmodium exported protein, unknown function | RSLAE | |
| PF3D7_0113000 | glutamic acid-rich protein | RLLNE | |
| PF3D7_0113200 | Plasmodium exported protein, unknown function | RILAD | iRBC cytosol, partially localised to Maurer's clefts |
| PF3D7_0113300 | Plasmodium exported protein (hyp1), unknown function | RLLTE | iRBC |
| PF3D7_0113400 | Plasmodium exported protein, unknown function | RVLTE | |
| PF3D7_0113700 | heat shock protein 40, type II | RCLAE | iRBC |
| PF3D7_0113900 | Plasmodium exported protein (hyp8), unknown function | RWLSE | Maurer's Clefts |
| PF3D7_0114000 | exported protein family 1 | RILYS | Maurer's Clefts |
| PF3D7_0201500 | Plasmodium exported protein (hyp9), unknown function | RSLSE | |
| PF3D7_0201600 | PHISTb domain-containing RESA-like protein 1 | RNLSS | iRBC cytosol, iRBC periphery |
| PF3D7_0201700 | DnaJ protein, putative | RQLSE | |
| PF3D7_0201800 | knob associated heat shock protein 40 | RNLAQ | PVM, iRBC cytosol |
| PF3D7_0201900 | erythrocyte membrane protein 3 | RSLAQ | iRBC periphery (Knobs) |
| PF3D7_0202000 | knob-associated histidine-rich protein | RTLAQ | iRBC periphery (Knobs) |
| PF3D7_0204100 | conserved Plasmodium protein, unknown function | | iRBC cytosol |
| PF3D7_0214000 | T-complex protein 1 subunit theta (CCT8) | | iRBC cytosol |
| PF3D7_0214100 | protein transport protein SEC31 (SEC31) | | Punctate signal in iRBC cytosol |
| PF3D7_0219800 | Plasmodium exported protein (PHISTc), unknown function | RNLAQ | |
| PF3D7_0219900 | Plasmodium exported protein, unknown function | RNLIE | |
| PF3D7_0220100 | DnaJ protein, putative | RSLCE | |
| PF3D7_0220300 | Plasmodium exported protein, unknown function | RTLTE | |
| PF3D7_0220500 | Plasmodium exported protein (hyp2), unknown function | RTLNY | |
| PF3D7_0220700 | Plasmodium exported protein (hyp9), unknown function | RLLVE | |
| PF3D7_0220800 | cytoadherence linked asexual protein 2 (CLAG2) | | PVM, iRBC periphery |
| PF3D7_0221700 | Plasmodium exported protein, unknown function | RILSD | |
| PF3D7_0222100 | Pfmc-2TM Maurer's cleft two transmembrane protein | RMLAQ | Maurer's Clefts |
| PF3D7_0301300 | alpha/beta hydrolase, putative | RYLSE | iRBC cytoskeleton |
| PF3D7_0301400 | Plasmodium exported protein, unknown function | RLLEE | |
| PF3D7_0301500 | Plasmodium exported protein, unknown function | RSLYE | |
| PF3D7_0301600 | Plasmodium exported protein (hyp1), unknown function | RLLAQ | iRBC cytosol |
| PF3D7_0301700 | Plasmodium exported protein, unknown function | | Maurer's Clefts |
| PF3D7_0302500 | cytoadherence linked asexual protein 3.1 (CLAG3.1) | | PVM, iRBC periphery |
| PF3D7_0310400 | parasite-infected erythrocyte surface protein | RLLVD | TVN junction protein, iRBC periphery, iRBC cytoplasm and around PV |
| PF3D7_0312400 | glycogen synthase kinase 3 (GSK3) | | Maurer's Clefts |
| PF3D7_0324100 | Pfmc-2TM Maurer's cleft two transmembrane protein | RILAQ | Maurer's Clefts |
| PF3D7_0401800 | Plasmodium exported protein (PHISTb), unknown function | RNLSE | iRBC periphery |
| PF3D7_0402000 | Plasmodium exported protein (PHISTa), unknown function | RNLSE | PVM |
| PF3D7_0402100 | Plasmodium exported protein (PHISTb), unknown function | RKLYHE | |
| PF3D7_0402400 | Plasmodium exported protein, unknown function | RILVE | J-dots |
| PF3D7_0404800 | conserved Plasmodium protein, unknown function | RVLHN | |
| PF3D7_0410000 | erythrocyte vesicle protein 1 (EVP1) | RIIAE | iRBC |
| PF3D7_0422000 | steroid dehydrogenase, putative | RDLLE | |
| PF3D7_0424000 | Plasmodium exported protein (PHISTc), unknown function | RNLSE | |
| PF3D7_0424400 | surface-associated interspersed protein 4.2 (SURFIN 4.2) (SURF4.2) | | Maurer's Clefts, iRBC cytoplasm |
| PF3D7_0424500 | serine/threonine protein kinase, FIKK family (FIKK4.1) | RHLTE | Maurer's Clefts |
| PF3D7_0424600 | Plasmodium exported protein (PHISTb), unknown function | RILSE | iRBC periphery |
| PF3D7_0424700 | serine/threonine protein kinase, FIKK family (FIKK4.2) | RNLSE | K-dots |
| PF3D7_0424800 | Plasmodium exported protein (PHISTb), unknown function | RILST | |
| PF3D7_0424900 | Plasmodium exported protein (PHISTa), unknown function | RNLVQ | |
| PF3D7_0425100 | Plasmodium exported protein (hyp6), unknown function | RTLTE | |
| PF3D7_0500800 | mature parasite-infected erythrocyte surface antigen | RILSE | iRBC periphery |
| PF3D7_0501000 | Plasmodium exported protein, unknown function | RVLAE | Maurer's clefts |
| PF3D7_0501200 | parasite-infected erythrocyte surface protein | RTLAD | iRBC surface |
| PF3D7_0501300 | skeleton-binding protein 1 (SBP1) | | Partial co-localisation to Maurer's clefts and partial co-localisation to the iRBC periphery (Knobs) |
| PF3D7_0515300 | phosphatidylinositol 3-kinase (PI3K) | | iRBC cytoplasm |
| PF3D7_0532300 | Plasmodium exported protein (PHISTb), unknown function | RNLCE | iRBC cytosol, partially localised to Maurer's clefts |
| PF3D7_0532400 | lysine-rich membrane-associated PHISTb protein | RKLCE | iRBC periphery (Knobs), Partial co-localisation with Maurer's Clefts |
| PF3D7_0532500 | Plasmodium exported protein, unknown function | RSLSD | |
| PF3D7_0532600 | Plasmodium exported protein, unknown function | RILKQ | iRBC cytosol |
| PF3D7_0601500 | Plasmodium exported protein (PHISTb), unknown function | RNLSE | |
| PF3D7_0628400 | conserved Plasmodium membrane protein, unknown function | RSLYF | |
| PF3D7_0701600 | Pfmc-2TM Maurer's cleft two transmembrane protein | RMLAQ | Maurer's Clefts |
| PF3D7_0701900 | Plasmodium exported protein, unknown function | RILIQ | |
| PF3D7_0702400 | small exported membrane protein 1 (SEMP1) | | Maurer's Clefts |
| PF3D7_0702500 | Plasmodium exported protein, unknown function | RILKS | Maurer's Clefts |
| PF3D7_0718100 | exported serine/threonine protein kinase (EST) | | iRBC periphery |
| PF3D7_0718300 | cysteine repeat modular protein 2 (CRMP2) | | Maurer's Clefts |
| PF3D7_0719800 | conserved protein, unknown function | KLIEE | |
| PF3D7_0721100 | conserved Plasmodium protein, unknown function | RKLLS | |
| PF3D7_0726100 | Plasmodium exported protein, unknown function | RSLYE | |
| PF3D7_0730900 | EMP1-trafficking protein | RSLTE | iRBC cytosol, partially localised to Maurer's clefts |
| PF3D7_0731100 | EMP1-trafficking protein | RNLGE | iRBC exosome-like vesicles |
| PF3D7_0731200 | Plasmodium exported protein, unknown function | RLLCE | |
| PF3D7_0731300 | Plasmodium exported protein (PHISTb), unknown function | RILSE | |
| PF3D7_0731700 | Plasmodium exported protein (hyp9), unknown function | RLLSQ | |
| PF3D7_0800800 | Plasmodium exported protein (hyp7), unknown function | RSLSS | |

| | | | |
|---|---|---|---|
| PF3D7_0801000 | Plasmodium exported protein (PHISTc), unknown function | RNLGA | J-dots |
| PF3D7_0807700 | serine protease DegP | RILND | iRBC cytosol |
| PF3D7_0814100 | conserved Plasmodium protein, unknown function | RFLTD | |
| PF3D7_0822600 | protein transport protein SEC23 (SEC23) | | Maurer's Clefts |
| PF3D7_0830400 | conserved Plasmodium protein, unknown function | | Maurer's Clefts |
| PF3D7_0830500 | sporozoite and liver stage tryptophan-rich protein, putative (TryThrA) | | Punctate signal in iRBC cytosol |
| PF3D7_0830600 | Plasmodium exported protein (PHISTc), unknown function | RILYE | |
| PF3D7_0830700 | Plasmodium exported protein (hyp9), unknown function | RSLAL | |
| PF3D7_0830900 | Plasmodium exported protein, unknown function | RLLIE | |
| PF3D7_0831200 | DnaJ protein, putative | RYLYA | |
| PF3D7_0831400 | Plasmodium exported protein, unknown function | RLLTE | |
| PF3D7_0831500 | Plasmodium exported protein (PHIST), unknown function | RNLSD | |
| PF3D7_0831600 | cytoadherence linked asexual protein 8 (CLAG8) | | PVM, iRBC periphery |
| PF3D7_0831800 | histidine-rich protein II | RLLHE | Maurer's Clefts, iRBC periphery |
| PF3D7_0901700 | Plasmodium exported protein (hyp5), unknown function | RCLTE | |
| PF3D7_0901800 | Plasmodium exported protein, unknown function | RSLAE | |
| PF3D7_0902000 | serine/threonine protein kinase, FIKK family (FIKK9.1) | RSLCA | Maurer's Clefts |
| PF3D7_0902200 | serine/threonine protein kinase, FIKK family (FIKK9.3) | RSLSV | Maurer's Clefts |
| PF3D7_0902300 | serine/threonine protein kinase, FIKK family (FIKK9.4) | RKLAE | Maurer's clefts |
| PF3D7_0902500 | serine/threonine protein kinase, FIKK family (FIKK9.6) | RYLSE | Maurer's Clefts |
| PF3D7_0904900 | copper-transporting ATPase (CuTP) | | Parasite and iRBC periphery |
| PF3D7_0905400 | high molecular weight rhoptry protein 3 (RhopH3) | | PVM and iRBC periphery |
| PF3D7_0911300 | cysteine repeat modular protein 1 (CRMP1) | | Maurer's Clefts |
| PF3D7_0911900 | Falstatin (Inhibitor of cysteine protease) | | PV and puctated staining in iRBC cytoplasm |
| PF3D7_0929400 | high molecular weight rhoptry protein 2 (RhopH2) | | PVM and iRBC periphery |
| PF3D7_1001400 | alpha/beta hydrolase, putative | RSLGE | iRBC cytosol |
| PF3D7_1001600 | alpha/beta hydrolase, putative | RKLAE | iRBC cytoskeleton |
| PF3D7_1001700 | Plasmodium exported protein (PHISTc), unknown function | KILCE | |
| PF3D7_1001800 | Plasmodium exported protein (PHISTc), unknown function | RTLSD | |
| PF3D7_1001900 | Plasmodium exported protein (hyp16), unknown function | RFLSE | Lumen of Maurer's Clefts |
| PF3D7_1002000 | Plasmodium exported protein (hyp2), unknown function | RLLAE | |
| PF3D7_1002100 | EMP1-trafficking protein | RLLSE | |
| PF3D7_1002300 | conserved Plasmodium protein, unknown function | RNLYI | |
| PF3D7_1007000 | conserved membrane protein, unknown function | RILSV | |
| PF3D7_1013700 | conserved Plasmodium protein, unknown function | RILTE | |
| PF3D7_1016300 | glycophorin binding protein | RILAE | PVM and RBC vesicles (MCs?) |
| PF3D7_1016400 | serine/threonine protein kinase, FIKK family (FIKK10.1) | RCLAE | Maurer's Clefts |
| PF3D7_1016500 | Plasmodium exported protein (PHISTc), unknown function | RILSE | |
| PF3D7_1016600 | Plasmodium exported protein (PHISTc), unknown function | RILSE | Punctate signal in iRBC cytosol |
| PF3D7_1016800 | Plasmodium exported protein (PHISTc), unknown function | RVLTE | |
| PF3D7_1016900 | early transcribed membrane protein 10.3 | RALKD | PV (Trophozoites, Gametocytes), iRBC (Schizonts) |
| PF3D7_1033200 | early transcribed membrane protein 10.2 | RNLIL | iRBC cytosol |
| PF3D7_1038700 | Plasmodium exported protein, unknown function | RLLGE | |
| PF3D7_1038800 | RESA-like protein with PHIST and DnaJ domains | RKLYS | |
| PF3D7_1039000 | serine/threonine protein kinase, FIKK family (FIKK10.2) | RHLSD | iRBC |
| PF3D7_1102300 | Plasmodium exported protein, unknown function | RILCE | |
| PF3D7_1102500 | Plasmodium exported protein (PHISTb), unknown function | RNLYE | iRBC surface |
| PF3D7_1102900 | Plasmodium exported protein (hyp11), unknown function | RILTE | |
| PF3D7_1106800 | pseudo-tyrosine kinase-like protein | | iRBC |
| PF3D7_1121300 | tyrosine kinase-like protein (TKL2) | | iRBC |
| PF3D7_1127000 | protein phosphatase, putative | RCLNY | |
| PF3D7_1148900 | Plasmodium exported protein, unknown function | | Partially exported in P. falciparum. Maurer's Clefts |
| PF3D7_1149000 | antigen 332, DBL-like protein (Pf332) | | Partially exported in P. falciparum. Maurer's Clefts |
| PF3D7_1149200 | ring-infected erythrocyte surface antigen (PHISTb) | RNLYCE | iRBC periphery |
| PF3D7_1149300 | serine/threonine protein kinase, FIKK family (FIKK11) | RILYE | |
| PF3D7_1149600 | DnaJ protein, putative | RNLSE | |
| PF3D7_1200800 | serine/threonine protein kinase, FIKK family (FIKK12) | KCLSE | Maurer's Clefts |
| PF3D7_1201000 | Plasmodium exported protein (PHISTb), unknown function | RILSS | PVM and iRBC periphery |
| PF3D7_1201100 | RESA-like protein with PHIST and DnaJ domains | RYLCE | |
| PF3D7_1201200 | Plasmodium exported protein (PHISTa-like), unknown function | RKLAD | |
| PF3D7_1212100 | peripheral plastid protein 1, putative | RILKE | |
| PF3D7_1227200 | potassium channel K1 | | iRBC periphery |
| PF3D7_1229400 | macrophage migration inhibitory factor | | iRBC |
| PF3D7_1237900 | conserved Plasmodium protein, unknown function | RNLCYE | |
| PF3D7_1244500 | conserved Plasmodium protein, unknown function | RDLIK | |
| PF3D7_1252300 | conserved Plasmodium protein, unknown function | | Maurer's Clefts |
| PF3D7_1252500 | Plasmodium exported protein, unknown function | RILEE | |
| PF3D7_1252700 | Plasmodium exported protein (PHISTb), unknown function | RTLFE | iRBC periphery |
| PF3D7_1252800 | Plasmodium exported protein (PHISTb), unknown function | RILCT | |
| PF3D7_1252900 | Plasmodium exported protein, unknown function | RTLVE | |
| PF3D7_1253000 | gametocyte erythrocyte cytosolic protein | RILSD | iRBC cytosol (gametocytes) |
| PF3D7_1253100 | Plasmodium exported protein (PHISTa), unknown function | RNLCE | |
| PF3D7_1302000 | EMP1-trafficking protein | RSLSE | |
| PF3D7_1310500 | conserved protein, unknown function | | PV, iRBC periphery |
| PF3D7_1334300 | MSP7-like protein (MSRP5) | | Partially exported in P. falciparum. Punctated signal in iRBC cytosol |
| PF3D7_1334500 | MSP7-like protein (MSRP6) | | Punctated staining in iRBC cytosol |
| PF3D7_1334600 | MSP7-like protein (MSRP3) | KLLEE | |
| PF3D7_1334700 | MSP7-like protein (MSRP7) | | iRBC cytosol |
| PF3D7_1352900 | Plasmodium exported protein, unknown function | RKLAE | |
| PF3D7_1353100 | Plasmodium exported protein, unknown function | RILTQ | |
| PF3D7_1353200 | MAHRP2 | | Maurer's Clefts tethers |
| PF3D7_1370300 | membrane associated histidine-rich protein (MAHRP1) | | Maurer's Clefts |
| PF3D7_1372000 | Plasmodium exported protein (PHISTa), unknown function | RNLSE | |
| PF3D7_1372100 | Plasmodium exported protein (PHISTb), unknown function | RSLLG | |

| | | | |
|---|---|---|---|
| PF3D7_1372200 | histidine-rich protein III | RLLHE | |
| PF3D7_1372300 | Plasmodium exported protein (PHIST), unknown function | RNLAQ | |
| PF3D7_1401100 | DnaJ protein, putative | RNLSE | |
| PF3D7_1401200 | Plasmodium exported protein, unknown function | RSLAE | iRBC cytosol, partially localised to Maurer's clefts and J-dots |
| PF3D7_1401300 | proline aminopeptidase | RILCD | iRBC |
| PF3D7_1401400 | early transcribed membrane protein 14.1 (ETRAMP14) | | Maurer's Clefts |
| PF3D7_1401600 | Plasmodium exported protein (PHISTb), unknown function | RSLYE | |
| PF3D7_1425900 | conserved Plasmodium protein, unknown function | RNLIE | |
| PF3D7_1429600 | conserved Plasmodium protein, unknown function | RLLCT | |
| PF3D7_1439000 | Copper Transporter | | iRBC periphery (Rings), parasite membrane (Schizonts) |
| PF3D7_1447700 | conserved Plasmodium protein, unknown function | RLLLF | |
| PF3D7_1448400 | ubiquitin-protein ligase, putative (HRD3) | | iRBC |
| PF3D7_1458300 | conserved Plasmodium protein, unknown function | RALTY | |
| PF3D7_1467000 | conserved Plasmodium membrane protein, unknown function | RLLEG | |
| PF3D7_1476200 | Plasmodium exported protein (PHISTb), unknown function | RCLSE | iRBC periphery |
| PF3D7_1476300 | Plasmodium exported protein (PHISTb), unknown function | RKLYE | |
| PF3D7_1477500 | Plasmodium exported protein (PHISTb), unknown function | RNLSD | iRBC cytosol, partially localised to Maurer's clefts |
| PF3D7_1478000 | Plasmodium exported protein (PHISTa), unknown function | RNLTE | |
| PF3D7_1478100 | Plasmodium exported protein (hyp13), unknown function | RSLAE | |
| PF3D7_1478600 | EMP1-trafficking protein | RSLYE | |
| PF3D7_1478800 | Plasmodium exported protein, unknown function | RRLSE | |
